# The effect of external stimulation on functional networks in the aging healthy human brain

**DOI:** 10.1101/2021.11.19.469206

**Authors:** Anira Escrichs, Yonatan Sanz Perl, Noelia Martínez-Molina, Carles Biarnes, Josep Garre-Olmo, José Manuel Fernández-Real, Rafel Ramos, Ruth Martí, Reinald Pamplona, Ramon Brugada, Joaquin Serena, Lluís Ramió-Torrentà, Gabriel Coll-De-Tuero, Luís Gallart, Jordi Barretina, Joan C. Vilanova, Jordi Mayneris-Perxachs, Luca Saba, Salvador Pedraza, Morten L. Kringelbach, Josep Puig, Gustavo Deco

## Abstract

Understanding the brain changes occurring during aging can provide new insights for developing treatments that alleviate or reverse cognitive decline. Neurostimulation techniques have emerged as potential treatments for brain disorders and to improve cognitive functions. Nevertheless, given the ethical restrictions of neurostimulation approaches, *in silico* perturbation protocols based on causal whole-brain models are fundamental to gaining a mechanistic understanding of brain dynamics. Furthermore, this strategy could serve as a more specific biomarker relating local activity with global brain dynamics. Here, we used a large resting-state fMRI dataset divided into middle-aged (N=310, aged < 65 years) and older adults (N=310, aged *≥* 65) to characterize brain states in each group as a probabilistic metastable substate (PMS) space, each with a probabilistic occurrence and frequency. Then, we fitted the PMS to a whole-brain model and applied *in silico* stimulations with different intensities in each node to force transitions from the brain states of the older group to the middle-age group. We found that the precuneus, a brain area belonging to the default mode network and the rich club, was the best stimulation target. These findings might have important implications for designing neurostimulation interventions to revert the effects of aging on whole-brain dynamics.

## Introduction

Normal aging causes changes in the brain that can lead to cognitive decline, thereby affecting the quality of life and autonomy of the elderly and their caregivers (Barnes, 2011; Li et al., 2015). Longitudinal studies in healthy older adults have shown an association between altered functional connectivity in resting-state and decreased cognitive functions (Fjell et al., 2017; Persson et al., 2014), thus suggesting that the resting-state could be an indicator of age-related cognitive decline. In addition, various neuroimaging studies have described that aging affects several resting-state networks (Betzel et al., 2014; Ferreira and Busatto, 2013; Grady et al., 2016; Spreng et al., 2016; Wang et al., 2010), and the rich-club organization of the human brain (Cao et al., 2014; Damoi-seaux, 2017; Escrichs et al., 2021a; Zhao et al., 2015). However, a question that remains to be addressed is whether these effects could be reversed or alleviated with external stimulation protocols that promote transitions from the brain states of the older towards those observed in younger adults.

The study of causal structure-function inferences has enhanced the understanding of the mechanisms underlying human brain dynamics, both through direct neurostimulation techniques (Casali et al., 2013; Ozdemir et al., 2020) and by *in silico* stimulation protocols (Bolton et al., 2020; Deco et al., 2018, 2019; Kringelbach and Deco, 2020; Muldoon et al., 2016). Non-invasive neurostimulation techniques such as transcranial electrical stimulation (tES) and transcranial magnetic stimulation (TMS) combined with neuroimaging have provided novel insights into the underlying mechanisms of stimulation-induced effects along with its impact on large-scale functional brain networks (Bestmann and Feredoes, 2013). These approaches have emerged as potential treatments for neurological and neurodegenerative disorders (Fox et al., 2014; Kunze et al., 2016) as well as for improving cognitive function in healthy individuals (Clark and Parasuraman, 2014). Nevertheless, experimental and ethical constraints limit the exploration of efficient practices that could be improved by the inclusion of whole-brain computational approaches along with *in silico* perturbations. In particular, dynamical models of brain activity have been fitted to different brain states to systematically apply *in silico* perturbations that promote transitions between brain states, and consequently, predict optimal neurostimulation targets (Deco et al., 2019; Ipiña et al., 2020; Muldoon et al., 2016). This strategy allows exploring dynamical brain responses elicited by controlled perturbative protocols, which are not constrained by ethical limitations (Deco et al., 2017).

In this context, we postulate that causal whole-brain modeling along with *in silico* stimulations can promote the transition between brain states of different age groups characterized by their dynamical behavior where the external stimulation represents the perturbation needed to induce that transition. The first step to finding support for this interpretation is to define the brain states associated with aging through their underlying dynamical behavior, thus providing a quantitative characterization. The probability metastable substates (PMS) space emerges as an optimal space to describe this dynamical behavior as the time evolution of a set of metastable states obtained within the Leading Eigenvector Dynamical Analysis (LEiDA) (Cabral et al., 2017; Deco et al., 2019; Kringelbach and Deco, 2020). The LEiDA framework has allowed discerning brain states in depression (Figueroa et al., 2019), different states of consciousness (Deco et al., 2019; Kringelbach et al., 2020; Kringelbach and Deco, 2020; Lord et al., 2019) and healthy aging (Cabral et al., 2017). Notably, in our previous work Escrichs et al. (2021a), we found significant differences in the PMS space between old and middle-aged healthy participants groups. The second step to support our hypothesis involves the transition from the older subjects’ PMS representation to the youngest one induced by *in silico* perturbations. This can be done through whole-brain models, which link the underlying anatomical connectivity with functional dynamics obtained from neuroimaging data, in which the external stimulation of all brain areas can be systematically explored via *in silico* perturbations by adjusting the parameters of the model (Deco et al., 2018, 2019; Kringelbach and Deco, 2020). In other words, the empirical LEiDA approach obtains the PMS of each group, while the model-based *in silico* approach allows us to simulate the PMS space of the older group and artificially perturb each brain area to induce transitions towards the PMS of the middle-age group. This mechanistic approach allows for an effective way of perturbing the model by simply changing the bifurcation parameter in a given brain area.

Here, we extended our previous publication (Escrichs et al., 2021a) for applying a causal mechanistic approach which forces transitions between brain states promoted by *in silico* perturbations (Deco et al., 2019; Kringelbach and Deco, 2020). We used a large resting-state neuroimaging dataset of healthy human adults divided into two groups: middle-age group (N=310, aged<65 years) and older group (N=310, aged*≥*65). First, we obtained the dynamical description by computing the probabilistic metastable substates (PMS) space over the empirical fMRI data of each group through the LEiDA framework. Second, we implemented a Hopf whole-brain model and optimized the model’s parameters to simulate the PMS space of the older group. Finally, we applied *in silico* perturbations to the optimized model of the older group to rebalance it toward the empirical PMS of the middle-age group as a reference of a healthy regime state.

## Materials and Methods

### Participants

Neuroimaging data were obtained from the Aging Imageomics Study (Puig et al., 2020) and comprised 620 healthy adults divided into two groups. The middle-aged group comprised 310 subjects aged < 65 years (mean age, 60.2*±*3.7 y), and the older group comprised 310 subjects aged >= 65 years (mean age, 71.8*±*4.5 y). The experimental protocol was approved by the Ethics Committee of the Dr. Josep Trueta University Hospital. Written informed consent was obtained from all participants. A complete description of the neuroimaging data can be consulted in Puig et al. (2020) and Escrichs et al. (2021a).

### Resting-state acquisition and preprocessing

Imaging was performed on a mobile 1.5T scanner (Vantage Elan, Toshiba Medical Systems) with an 8-channel phased-array head coil with foam padding and headphones to restrict head motion and scanner noise. The high-resolution T1-weighted images were acquired with 112 slices in the axial plane (repetition time (TR) = 8 ms; echo time (TE) = 4.5 ms; flip angle = 15°; field of view (FOV) = 235 mm; and voxel size = 1.3×1.3×2.5 mm). Rs-fMRI scans were acquired axially for 5 minutes using a gradient echoplanar imaging sequence (122 volumes; TR = 2500 ms; TE = 40 ms; flip angle = 83°; FOV = 230 mm; voxel size = 3.5×3.5×5 mm; no gap). Participants were asked to remain motionless as possible and close their eyes.

T1 and EPI images were automatically oriented using Conn (Whitfield-Gabrieli and Nieto-Castanon, 2012). Processing Assistant for Resting-State fMRI (DPARSF) [(Chao-Gan and Yu-Feng, 2010), www.rfmri.org/DPARSF], which is based on Statistical Parametric Mapping (SPM12) (http://www.fil.ion.ucl.ac.uk/spm) was used to preprocess the MRI data. Preprocessing steps included: discarding the first 5 volumes from each scan to allow for signal stabilization; slice-timing correction; realignment for head motion correction across different volumes; T1 co-registration to the functional image; European regularisation segmentation; removal of spurious variance through linear regression: six parameters from the head motion correction, the white matter (WM) signal, and the cerebrospinal fluid signal (CSF) using CompCor (Behzadi et al., 2007); removal of the linear trend; spatial normalization to the Montreal Neurological Institute (MNI) standard space; spatial smoothing with 6 mm FWHM Gaussian Kernel; and band-pass temporal filtering (0.01-0.020 Hz). Finally, the time series for each subject were extracted using a resting-state atlas of 214 nodes (Shen et al., 2013).

### Difussion Tensor Imaging (DTI) acquisition and preprocessing

For the whole-brain model, we used an average structural connectivity matrix (SC) from a sample of 38 unrelated healthy subjects previously described in De Filippi et al. (2021). MRI images were acquired on a 3T whole-body Siemens TRIO scanner (Hospital Clínic, Barcelona) using a dual spin-echo DTI sequence (TR = 680ms; TE = 92ms; FOV = 236mm; 60 contiguous axial slices; isotropic voxel size 2×2×2 mm; no gap, and 118 ×118 matrix sizes). Diffusion was obtained with 64 optimal noncollinear diffusion directions using a single b value = 1,500s/mm2 interleaved with 9 non-diffusion b0 images. A frequency-selective fat saturation pulse was used to avoid chemical shift misregistration artifacts.

The whole-brain structural connectivity matrix (SC) was computed following the procedure applied in previous studies (Cao et al., 2013; Gong et al., 2009; López-González et al., 2021; Muthuraman et al., 2016). For each subject, a 214×214 SC was computed using the processing pipeline of the FMRIB’s Diffusion Toolbox (FDT) in FMRIB’s Software Library www.fmrib.ox.ac.uk/fsl. Non-brain tissues were extracted with Brain Extraction Tool (BET) (Smith, 2002), eddy current distortions and head motion were corrected using eddy correct (Andersson and Sotiropoulos, 2016), and the gradient matrix was reoriented to correct for subject motion (Leemans and Jones, 2009). Crossing fibres were modeled using BEDPOSTX, and the probability of multi-fibre orientations was computed to improve the sensitivity of non-dominant fibre populations (Behrens et al., 2003, 2007). The probabilistic tractography analysis was performed for each participant in native diffusion space using PROBTRACKX. The connectivity probability *SC*_*np*_ between brain areas *n* and *p* was calculated as the total proportion of sampled fibres in all voxels in brain area *n* that reach any voxel in brain area *p*. The *SC*_*np*_ matrix was then symmetrized by computing their transpose matrix *SC*_*pn*_ and averaging both matrices. Finally, averaging the resulting matrices across all participants, a whole-brain SC DTI matrix was obtained, representing a template of healthy adults.

### LEiDA

We characterized the empirical brain states by applying the Leading Eigenvector Dynamics Analysis (LEiDA) (Cabral et al., 2017; Deco et al., 2019; Kringelbach and Deco, 2020). This analysis was described in our previous study using the same fMRI dataset (Escrichs et al., 2021a). In particular, we extracted the time series for each participant using a resting-state atlas of 214 brain areas and computed the Hilbert transform to obtain the phase of the BOLD signals in every time-point for all brain areas of the parcellation **(Figure 1.A)**. Then, we computed a dynamic phase coherence connectivity matrix with size *N×N×T*, where *N=214* is the total brain areas, and *T=117* the total time-points. The BOLD phase coherence matrix **(Figure 1.B)** in each time *t* between each pair of brain areas *n* and *p* was estimated by computing the cosine of the phase difference as:

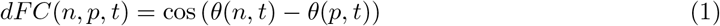

**Figure 1:**
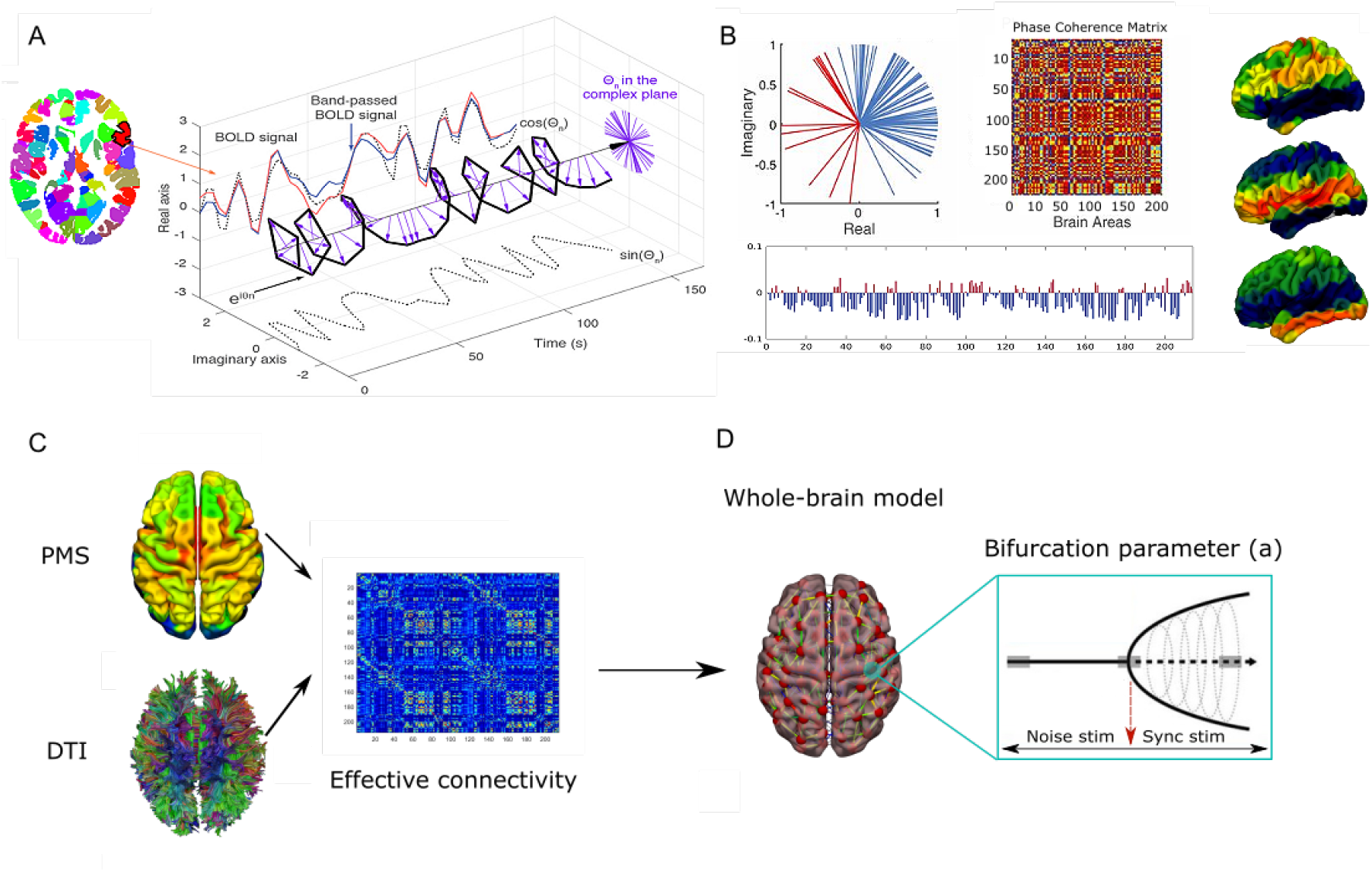
Probabilistic metastable substate (PMS) space, optimizing the model for whole-brain activity and *in silico* stimulations. (A) Time series extraction for each subject, brain area, and time point using the Hilbert transform. The panel shows a complex plane representing the BOLD phases for a given brain area across time. (B) Leading Eigenvector Dynamic Analysis (LEiDA) to identify dynamic functional connectivity patterns across all subjects [i.e., probabilistic metastable substates (PMS)]. The left panel shows the BOLD phases in all 214 brain areas described in the complex plane. The right panel shows the phase coherence matrix between each pair of brain areas in all time points. The vector shows the leading eigenvector *V*_1_(t), capturing the principal orientation of the BOLD phase (showing positive or negative values) for each of the 214 brain areas. The clustering configuration that best represented our resting-state fMRI data was found for 3 states. Rendered brains show the brain states rendered onto the cortex. (C) Whole-brain PMS model. A whole-brain dynamical model was fitted for the PMS of the older group based on the effective connectivity. (D) Stimulations *in silico*. Each brain area of the whole-brain model was systematically perturbed via *in silico* stimulations through two different protocols (noise and synchronization). The noise protocol shifts the local bifurcation parameter of each brain area to negative values, whereas the synchronization protocol shifts it to positive.

Given that the Hilbert Transform expresses any signal in the polar coordinate system (i.e., *x*_*a*_(*t*) = *A*(*t*) *·* cos (*φ*(*t*))), applying the cosine function to brain areas *n* and *p* with similar angles at a given time *t* will show a phase coherence close to 1 (i.e., cos(0°)=1), whereas brain areas showing orthogonality will show a phase coherence near zero (i.e., cos(90°) = 0) (Deco et al., 2019). Second, to characterize the dFC patterns across all subjects and time-points, we obtained a leading eigenvector *V*_1_(t) for each dFC(*t*) at time *t* by capturing the dominant functional connectivity pattern rather than the whole matrices. This approach allows to reduce dimensionality on the data considerably given that only considers a *V*_1_(t) for each dynamic FC matrix. The *V*_1_(t) is a Nx1 vector capturing the principal orientation of the BOLD phase (showing positive or negative values) for each of the 214 brain areas **(Figure 1.B lower panel)**. Next, we applied a k-means clustering algorithm ranged from *k* = 2 to *k* = 7 clusters to detect metastable substates or dynamic FC states from all the leading eigenvectors *V*_1_(*t*) across time-points, number of subjects, and groups to identify recurrent dynamic FC patterns across subjects. The total of leading eigenvectors were 117 time-points × 310 subjects × 2 groups = 72,540 *V*_1_(*t*). We obtained k cluster centroids, each one as an N×1 vector representing recurrent metastable substates across all participants. The resulting k-cluster centroids define the metastable substates among which the brain dynamics are switching across time, and the probability of occurrence of each substate determines the PMS of the brain. As a proof-of-concept, we display the minimum number of clusters that statistically differed between groups. The clustering configuration that best represented the resting-state data across all participants and distinguished between both groups was detected at k=3. **Figure 1.B right panel** shows the cluster centroid vectors onto the surface cortex using Surf Ice (https://www.nitrc.org/projects/surfice/).

### Whole-Brain Computational Model

The whole-brain BOLD activity was simulated using the so-called Hopf computational model, linking the anatomy and function. The model consisted of 214 dynamical cortical and subcortical brain areas coupled with the SC matrix. The local dynamics of each brain area was described by the normal form of a supercritical Hopf bifurcation, which emulates the dynamics for each brain area from noisy to oscillatory dynamics as follows:

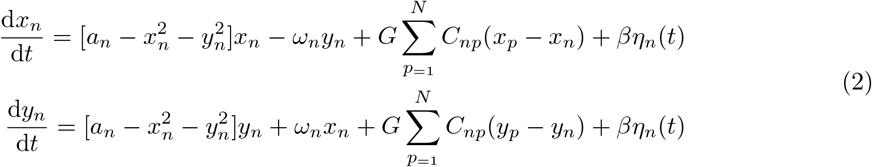

where *η*_*n*_(*t*) is additive Gaussian noise with standard deviation *β* = 0.02, and *C*_*np*_ is the SC that couples the local dynamics of brain area *n* with *p* and was normalized to a maximum value of *C* = 0.2. This normal form has a supercritical bifurcation at *a*_*n*_ = 0, such that for *a*_*n*_ > 0 the system is in a stable limit cycle oscillation with frequency *f*_*n*_ = *ω*_*n*_/2*π*, whereas for *a*_*n*_ < 0 the local dynamics are in a stable point (i.e., noisy state). The frequency *ω*_*n*_ of each brain area was estimated from the data which was given by the applied narrowband (i.e., 0.04 *−* 0.07*Hz*). The variables *x*_*n*_ emulate the BOLD signal of each node *j*. The global coupling factor G (scaled equally for each brain area) is the control parameter which allows adjusting the model to obtain the optimal dynamical working point where the simulations maximally fit the empirical data. We simulated the PMS as a function of the global coupling parameter G through the underlying structural connectivity matrix. We improved the fitting of the whole-brain model through the inclusion of the effective connectivity (EC) **(Figure 1.C)**, where the anatomical connectivity was updated by the synaptic weights that take into account the empirical functional connectivity. The effective connections were computed by measuring the distance between the empirical 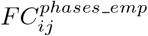 and the model 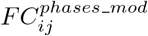 grand-averaged phase coherence matrices, and adjusted each structural connection *ij* separately using a gradient-descent approach. The model initially started computing with the SC matrix obtained from DTI and was run repeatedly with the updated EC matrix until the fit converged toward a stable value using the following procedure:

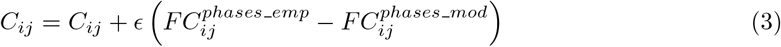

where *ϵ* = 0.01, and the grand average phase coherence matrices were defined as:

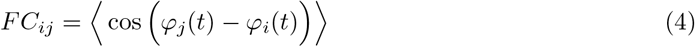

where *φ*(*t*) corresponds to the BOLD signal phase (obtained by the Hilbert transform) of the brain areas *j* and *i* at time *t*, and the brackets correspond to the average across time.

Finally, we systematically perturbed the 214 brain areas of the whole-brain model through two different protocols (noise and synchronization), which were based on shifting the local bifurcation parameter (*a*) of the optimized model **(Figure 1.D)**. The noise protocol applies negative (positive) intensities to the local parameter from 0 to *−*0.3 (0.1).

### Comparing empirical and simulated probability metastable space states

The empirical and simulated brain states were compared by using a symmetrized KL distance between the simulated and empirical probabilities as:

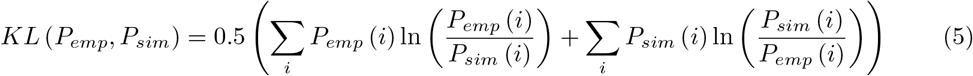

where *P*_*emp*_(*i*) are the empirical and *P*_*sim*_(*i*) the simulated probabilities on the same empirical extracted brain states *i*. The optimal simulated PMS is defined by the minimum KL distance between the empirical and simulated PMS.

## Results

### Leading Eigenvector Dynamics Analysis (LEiDA)

**Figure 2** left panel shows the probability of occurrence for the PMS of each group. The probability of the first metastable substate occurrence was higher in the older group than in the middle-age group [0.476 ± 0.008 (mean ± SE) vs. 0.453 ± 0.008 FDR-corrected p=0.03]. By contrast, the second metastable substate’s probability was higher in the middle-age group [0.288 ± 0.007 vs. 0.269 ± 0.006 in the older group, FDR-corrected = 0.026]. This state overlaps with the brain’s rich-club organization (i.e., the superior frontal cortex, precuneus, insula, and subcortical areas, such as the caudate, putamen, hippocampus, and thalamus), and its lower probability of occurrence in the older group can be interpreted as an alteration in the intrinsic dynamics within the rich-club or damage in any of their brain areas involved (Escrichs et al., 2021a). Consequently, we aimed to rebalance the PMS of the older group toward a younger regime (middle-aged group) through *in silico* stimulations.

**Figure 2:**
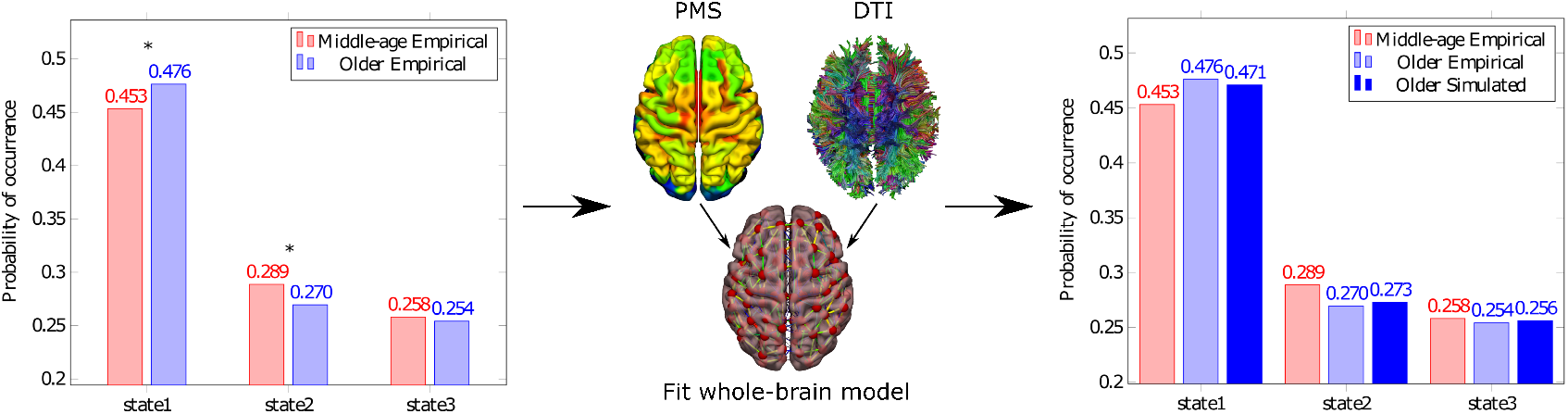
Empirical PMS and whole-brain model fitting. The left panel shows the empirical PMS of each group obtained by LEiDA. In red the middle-aged group and blue the older group. The probability of occurrence in the first state was higher in the older group. By contrast, the probability of occurrence in the second state was higher in the middle-age group. The whole-brain brain model of the older group was fitted to the empirical PMS of the older group. The right panel shows the resulting model (electric blue), remarkably similar to their empirical version (blue).

### Fit whole-brain computational model to the brain states of the older group

For the older group, we constructed a dynamical model of 214 non-linear oscillators representing the macroscopic dynamical behavior of each brain area of interest (**Figure 2** middle panel). These oscillators are coupled by a structural connectivity matrix (SC) among brain areas giving rise to collective dynamics. The local dynamics of each brain area was described by the normal form of a supercritical Hopf bifurcation, and the bifurcation parameters of each oscillator (*a*) were set in the edge of the bifurcation point, that is, the optimal point to represent the metastability of brain states (Deco et al., 2017). The coupling strength parameter (*G*) was optimized to fit the whole-brain model to the PMS of the older group. We used the centroids of the empirical PMS and built the model based on the probability of the empirical centers. Then, we estimated the distance between the model and the empirical phase coherence matrices and adjusted each structural connection separately using a gradient-descent algorithm. The model was run repeatedly with the updated EC until convergence to a stable point. We tested the differences between the empirical and the simulated probabilities by computing the symmetrized Kullback–Leibler (KL) distance. The optimal working point of the model was found at G=0.02, i.e., where the model fits the empirical PMS data of the older group. **Figure 2** right panel shows the generated model that reached an excellent fit between the empirical and the simulated probabilities (electric blue).

### *In silico* stimulations to force transitions between brain states

**Figure 3.A** shows the procedure to force transitions between brain states. We started from the simulated PMS that presented higher similarity to the empirical PMS of the older group (left panel, electric blue) and perturbed the model to force the transition to the empirical PMS of the middle-age group. To do so, we applied two different stimulation protocols: noise and synchronization, based on systematically shifting the local bifurcation parameter (*a*) of the optimized whole-brain model (middle panel). The noise protocol applies negative intensities to the local parameter, whereas the synchronization applies positive intensities. The strength of the perturbation is linked to the shifting of the local bifurcation parameter. We systematically perturbed each of the 214 brain areas of the whole-brain model and compared the distributions with the empirical PMS of the middle-age group. The optimal perturbation is that yields that the first brain state decreases, the second increases, and the third brain state remains similar (right panel).

**Figure 3:**
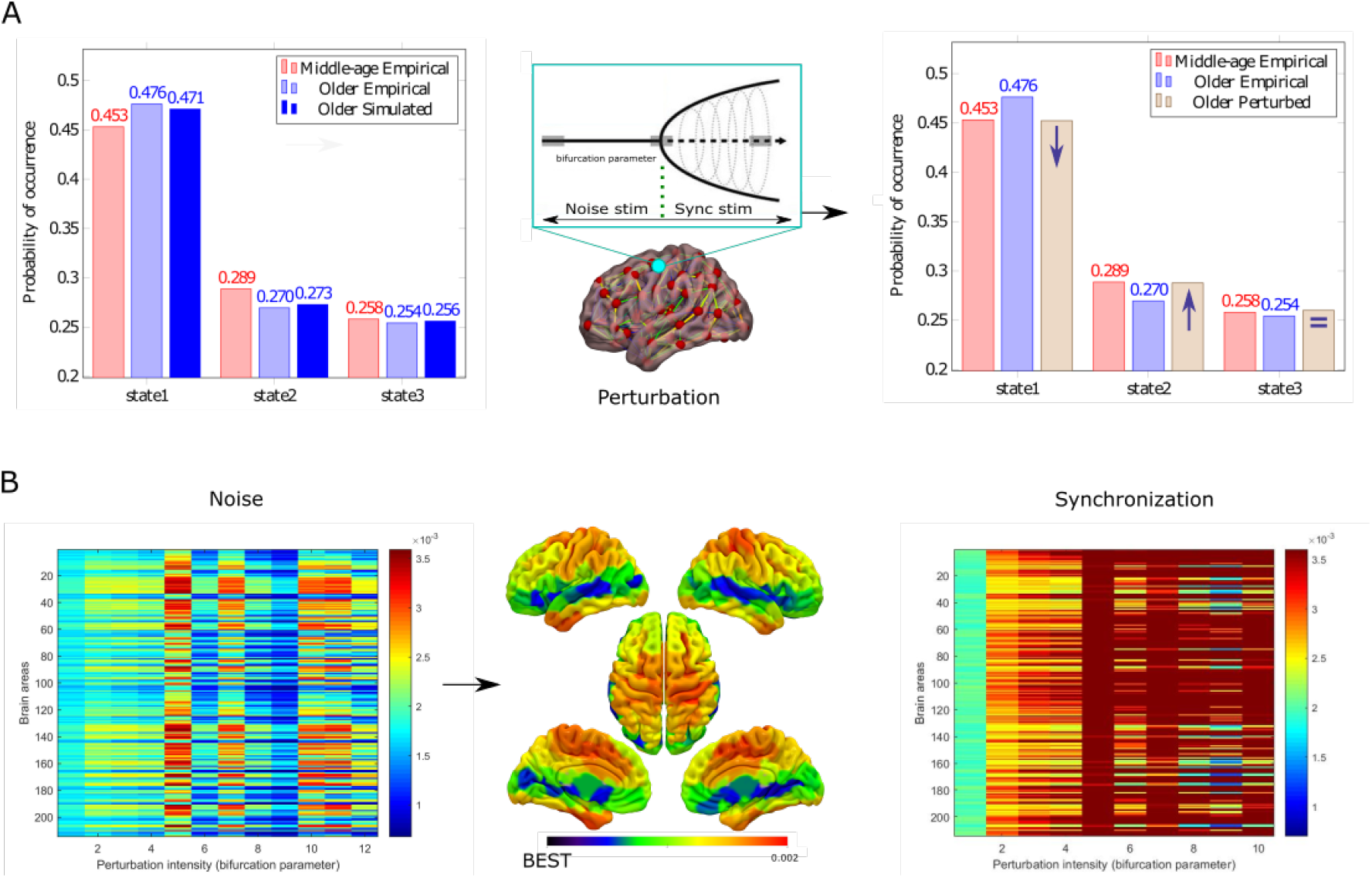
Noise and synchronization stimulation protocols. (A) Forcing transitions from the model of the older group (electric blue) to the empirical PMS of the middle-age group (red). The whole-brain model was perturbed at the optimal working point using two different protocols (noise and synchronization), which shifted the local bifurcation parameter to negative and positive values, respectively (middle panel). The optimal perturbation is that which achieves that the first brain state decreases, the second increases, and the third brain state remains similar (right panel). (B) The left matrix shows the KL-distance value after applying the noise protocol’s perturbation intensity (from softer to stronger) in each brain area. The brain rendered onto the cortex represents the KL-distance for the noise protocol between the PMS of the middle-age group and the perturbed model. In blue, potential brain areas to perturb to achieve a good transition between brain states. The right matrix shows that the synchronization protocol presented poor effectiveness given that KL distances were longer than in the noise protocol.

**Figure 3.B** shows the obtained matrices for each protocol, i.e., noise (left panel) and synchronization (right panel). The color scale represents the KL distance between the PMS of the middle-age group and the perturbed model. For the noise protocol, the KL distances were minimal in some brain areas, and thus a good transition between brain states was obtained. In contrast, the KL distances were higher in the synchronization protocol for all perturbation strengths and perturbed brain areas. This result indicates the unsuitability of the synchronization protocol to force the transition. The rendered brain shows that the precuneus, bilateral middle temporal gyrus, bilateral calcarine sulcus, bilateral inferior gyrus orbitofrontal part, left superior temporal gyrus, left insula, bilateral putamen, bilateral thalamus, and right caudate stand for those regions suitable to induce the transitions (middle panel).

**Figure 4** displays the comparison between the empirical PMS of each group, the optimized model of the older group, the model after perturbing the right precuneus with the best perturbation strength (*a* = *−*0.2250), and the model with the highest KL distance (the worst target). It is noticeable that for the right precuneus, the empirical and perturbed probability is almost the same for the three metastable substates considered. These results suggest that the right precuneus is the brain area that induces the best effective transition between brain states.

**Figure 4:**
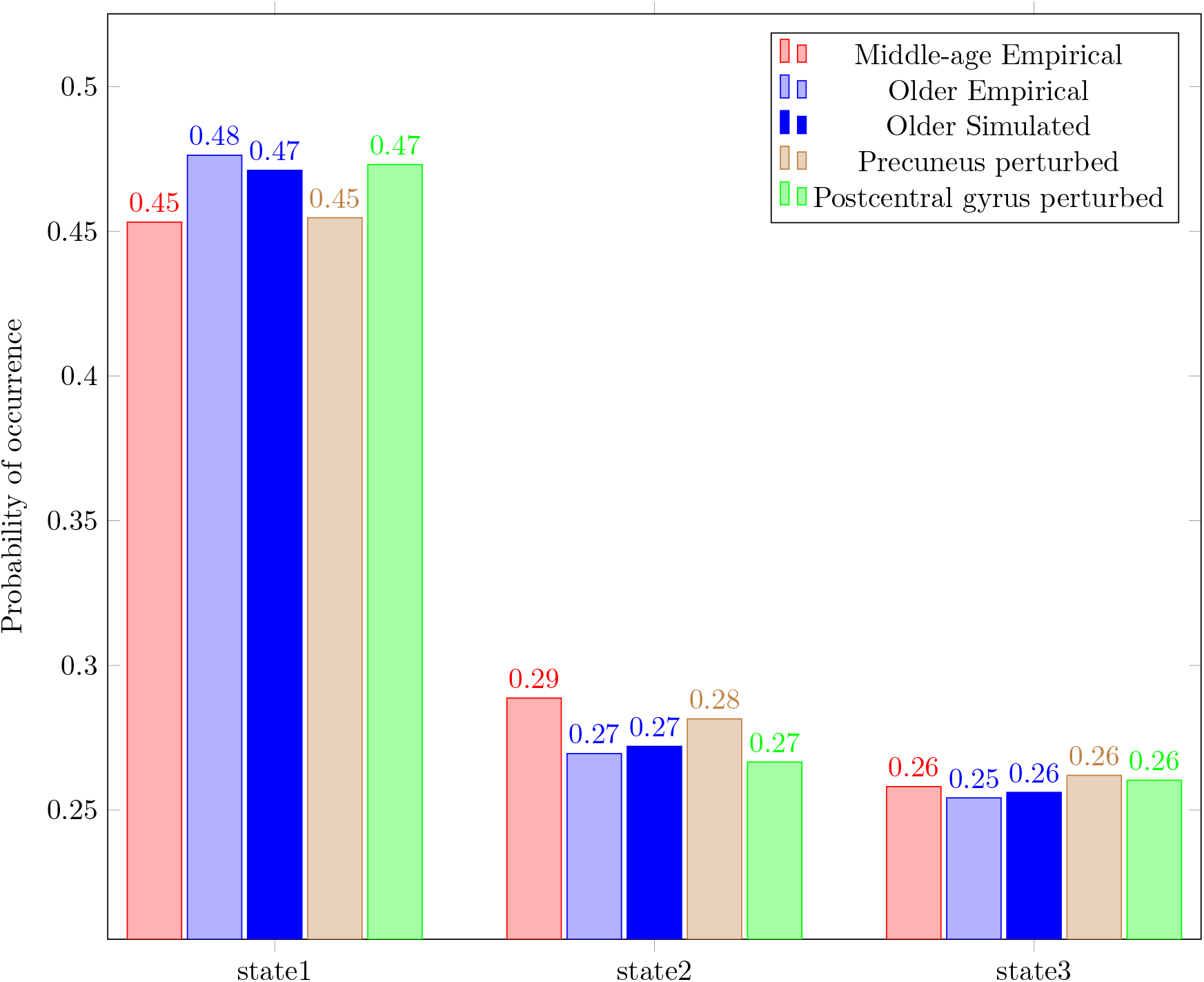
PMS comparison between empirical, modeled, and perturbed conditions. The plot shows the probability of occurrence of the empirical PMS of the middle-age group (in red) and the older group (in light blue). Electric blue represents the optimal fit for the PMS model of the older group. In brown, the model after changing the bifurcation parameter of the right precuneus using the noise protocol. This result indicates an optimal transition to the empirical PMS of the middle-age group (in red). This perturbation decreased the probability of occurrence of the first state, increased the probability of the second, and kept the probability of the third similar. The light green bars represent a non-optimal transition when the postcentral gyrus is stimulated.

## Discussion

In this work, we used empirical and computational approaches to study the causal dynamical mechanisms allowing the transition between brain states of different age groups. The empirical approach identified that older subjects have a lower probability of accessing a metastable substate that overlaps with the rich club. Such a state plays an essential role in integrating information across the whole-brain network and could explain brain dynamics alterations in the aging process. Then, we investigated the effect of perturbing all brain areas to induce optimal transitions from the states of the older aged group to the states of the middle-aged group by using causal whole-brain modeling and *in silico* perturbations. These results illustrated that forcing a shift in the intrinsic local dynamics of the right precuneus and other brain areas belonging to the rich club (insula, putamen, caudate, and thalamus) is suitable for inducing those transitions. Crucially, our model-based *in silico* approach provides causal evidence that external stimulations in specific local brain areas can reshape whole-brain dynamics in the aging brain. Importantly, this could provide new insights into the differential sensitivity of each brain area to *in silico* perturbations as a specific model-based biomarker relating local activity with global brain dynamics.

Understanding the underlying brain changes occurring during normal aging can contribute to developing treatments to reverse cognitive impairment. In this regard, non-invasive neurostimulation therapies stand as a promising intervention for brain disorders (Clark and Parasuraman, 2014; Kunze et al., 2016). Nevertheless, there are two different but related limitations for the application of such treatments. The first refers to the lack of a consensual definition of a brain state capable of being quantitatively characterized that differentiates the activity of an older from a younger brain. The second issue concerns the limitations to exploring the vast space of possible interventions due to experimental and ethical constraints (Deco et al., 2017). We addressed these two issues by applying whole-brain computational models, which allowed us to systematically explore brain responses elicited by *in silico* perturbations of fMRI empirical data of healthy older and middle-aged subjects.

Several attempts have been made to define and characterize the underlying dynamics of a given brain state (Escrichs et al., 2021b; Goldman et al., 2019; Kringelbach and Deco, 2020). Neuroimaging fMRI studies have motivated definitions related to static observables such as the functional connectivity (FC) or changes in the brain activity of RSNs associated with specific brain states (Alkire et al., 2008; Boly et al., 2012; Brodbeck et al., 2012; Tagliazucchi et al., 2013a,b; Vanhaudenhuyse et al., 2010). However, the dynamical behavior of brain activity, which is reflected in neuroimaging data, is often underlooked in these definitions. Recent works successfully defined the brain states through the probability of occurrence of metastable substates (PMS) space extracted from empirical fMRI data (Cabral et al., 2017; Deco et al., 2019). This definition captures the dynamical behavior of the states and presents high spatial overlap with RSNs (Deco et al., 2019; Figueroa et al., 2019; Kringelbach et al., 2020; Kringelbach and Deco, 2020; Lord et al., 2019). In particular, as shown here and in our previous work (Escrichs et al., 2021a), the empirical PMS analysis identified differences between older and middle-age groups in a metastable substate that closely overlaps with the so-called rich club (Hagmann et al., 2008; Sporns, 2013; van den Heuvel and Sporns, 2011), that in turn, the rich club overlaps with the DMN (Damoiseaux, 2017; van den Heuvel and Sporns, 2011). Our results reveal that, compared to middle-aged subjects, older subjects showed a lower probability of occurrence of this state, and when it occurred, it did so for shorter periods of time. In line with this finding, recent studies have suggested that the alterations in brain dynamics observed in the elderly could be due to a deficiency in the rich-club organization (Cao et al., 2014; Damoiseaux, 2017; Zhao et al., 2015).

We tested the hypothesis that causal modeling could predict optimal stimulation targets to rebalance the underlying brain dynamics in the elderly. Previous experimental studies investigated the effects of localized external perturbations during states of reduced awareness in humans (Angelakis et al., 2014; Thibaut et al., 2014; Zhang et al., 2019) and mild cognitive impairment (Hampstead et al., 2017). However, the systematic exploration via perturbations of all brain areas of the human brain can only be performed through computational models (Spiegler et al., 2016) that simulate the underlying brain activity. In this direction, recent works have implemented whole-brain models and *in silico* perturbations to explore the elicited responses from external stimulations in different brain states such as sleep, anesthesia, disorders of consciousness, and even in altered states such as meditation or the psychedelic state (Deco et al., 2019; Escrichs et al., 2021b; Ipiña et al., 2020; Kringelbach et al., 2020; Perl et al., 2020). In addition, perturbative models have been applied for individualized treatment in depressive (Ho et al., 2014) and epilepsy patients (Huang et al., 2017). Here, we modeled the PMS of the older group using a whole-brain computational model and applied artificial stimulations to force transitions between brain states of different ages. Interestingly, we show that this approach can be used to predict optimal targets to force transitions between brain states of different ages.

Specifically, our model-based *in silico* approach allowed us to test the effectiveness of two different stimulation protocols named noise and synchronization. The first protocol consists of reducing the value of the bifurcation parameter of the stimulated node resulting in noise outweighing oscillatory behavior, while the synchronization protocol yields the opposite effect. The fact that the noise protocol leads to better results means that the local bifurcation parameters must be mostly below or at the edge of bifurcation, thus favoring the local dynamics in the most susceptible regime. The results show that the brain area that promoted the best transition between brain states was the precuneus. The precuneus plays a central functional role in the DMN (Utevsky et al., 2014) and is involved in complex functions like memory, perception, mental imagery, and responses to pain (Cavanna and Trimble, 2006).

Furthermore, we found that the other brain areas that promoted an excellent transition are part of the so-called rich club (i.e., the precuneus, insula, putamen, caudate, and thalamus) (van den Heuvel et al., 2012; van den Heuvel and Sporns, 2011). Evidence suggests that a disruption in one of these regions can affect network efficiency and global brain function (van den Heuvel and Sporns, 2011). Rich-club regions are densely structurally interconnected, acting as a link for different functional modules in the brain, contributing to efficient communication and integration across the whole-brain network (van den Heuvel and Sporns, 2011). Additionally, considering that the protocol that achieved the best transition was the noise protocol, it might be related to brain overactivation that has been largely documented in the elderly. Particularly, older adults show overactivation in frontal brain areas (Cabeza et al., 2018; Davis et al., 2008; Reuter-Lorenz and Cappell, 2008; Yao and Hsieh, 2021), and among resting-state networks (Betzel et al., 2014; Escrichs et al., 2021a; Geerligs et al., 2015; Spreng et al., 2016). Thus, one possibility could be that noise stimulation decreases these functional overactivations.

Lastly, we would like to acknowledge some limitations in the study. One inherent limitation is related to using a cross-sectional approach that, by definition, cannot measure individual changes in brain dynamics. The image acquisition protocol with TR also limits this work = 2.5s in a 1.5T scanner. A protocol with increased spatial and temporal resolution could allow a more accurate representation of the underlying brain dynamics. Another limitation concerns the parcellation used, which was based on an atlas of 214 nodes. Using brain atlases with a large number of nodes could produce results with better local sensitivity.

Overall, the model-based *in silico* approach provides causal evidence that external stimulations in specific local brain areas can reshape whole-brain dynamics in aging. From a clinical standpoint, the methods and results presented here suggest optimal targets for neurostimulation techniques to induce transitions towards a healthy regime. This framework could improve the diagnosis, prognosis, and therapeutic responsiveness of aging effects in healthy adults and other conditions such as neuropsychiatry diseases and disorders of consciousness.

## Acknowledgments

AE, YS and GD were supported by the HBP SGA3 Human Brain Project Specific Grant Agreement 3 (grant agreement no. 945539), funded by the EU H2020 FET Flagship.

